# Genomic database furnishes a spontaneous example of a functional Class II glycyl-tRNA synthetase urzyme

**DOI:** 10.1101/2024.01.11.575260

**Authors:** Sourav Kumar Patra, Jordan Douglas, Peter R. Wills, Remco Bouckeart, Laurie Betts, Tang Guo Qing, Charles W. Carter

## Abstract

The chief barrier to studies of how genetic coding emerged is the lack of experimental models for ancestral aminoacyl-tRNA synthetases (AARS). We hypothesized that conserved core catalytic sites could represent such ancestors. That hypothesis enabled engineering functional “urzymes” from TrpRS, LeuRS, and HisRS. We describe here a fourth urzyme, GlyCA, detected in an open reading frame from the genomic record of the arctic fox, *Vulpes lagopus*. GlyCA is homologous to a bacterial heterotetrameric Class II GlyRS-B. Alphafold2 predicted that the N-terminal 81 amino acids would adopt a 3D structure nearly identical to the HisRS urzyme (HisCA1). We expressed and purified that N-terminal segment. Enzymatic characterization revealed a robust single-turnover burst size and a catalytic rate for ATP consumption well in excess of that previously published for HisCA1. Time-dependent aminoacylation of tRNA^Gly^ proceeds at a rate consistent with that observed for amino acid activation. In fact, GlyCA is actually 35 times more active in glycine activation by ATP than the full-length GlyRS-B α-subunit dimer. ATP-dependent activation of the 20 canonical amino acids favors Class II amino acids that complement those favored by HisCA and LeuAC. These properties reinforce the notion that urzymes represent the requisite ancestral catalytic activities to implement a reduced genetic coding alphabet.

## Introduction

Aminoacyl-tRNA synthetases, AARS, are the nanomachines that drive translation of genetic messages into proteins^1^. They, together with their cognate tRNAs, are the repository for all of the stereochemical information necessary to convert the strings of codons in genes into the alphabet of proteins. In that sense, they are “code-keys” kept by Nature’s locksmith. They also function as computational AND gates, synthesizing code-specific aminoacyl-tRNA molecules if and only if, within some appropriate tolerance, they bind simultaneously to the correct amino acid and the correct tRNA^2^. Their evolutionary history is central to resolving the origins of the intrinsic reflexivity—enforcing coding rules by which they assembled themselves—that forms the central challenge to understanding how genetic coding arose^3-5^. Experimental models for ancestral AARS•tRNA cognate pairs are thus vital to that pursuit because they provide the means to propose and test alternative evolutionary routes to the assembly of the coding table.

AARS urzymes^6-12^ are the most extensively studied such models. Ribozymes have been described that can either activate amino acids^13^, or acylate tRNAs with pre-activated amino acids^14, 15^. Unlike these ribozymes, the proteinaceous forms catalyze both amino acid activation and tRNA aminoacylation. Their catalytic rate accelerations exceed those of ribozymes by orders of magnitude. Moreover, they also exist in two Classes that discriminate appropriately between amino acids from their own class^2, 9, 16, 17^. These three properties give AARS urzymes special biological relevance.

Previously studied urzymes were all engineered for the purpose of testing the Rodin-Ohno^18^ hypothesis of bidirectional ancestry for Class I and II AARS. We constructed them on the basis of 3D superposition, truncating segments that were not highly conserved across the entire superfamily^12^. Thus, they were hypothesis-driven constructs and not driven by observations from natural sources.

We recently identified a sequence in GenBank whose 3D structure predicted by AlphaFold2^19^ closely resembles a Class II urzyme. This curious protein is a truncation of the α chain of the bacterial “orphan” glycyl-tRNA synthetase (GlyRS-B)^20^. It is purported to reside in an Arctic Fox (*Vulpes lagopus*) genome. However, it is more likely a bacterial contaminant. Only the gene for the α-subunit is present.

GlyRS-B is an α_2_β_2_ tetrameric Class II AARS found in bacteria and chloroplasts^21^. It is distinct from the homodimeric GlyRS-A found in archaea, eukaryotes, and some bacteria^3, 22^. GlyRS-B usually occurs as an α_2_β_2_ heterotetramer, with glycine activation being performed by the short α chains. Interestingly, in some organisms GlyRS-B is expressed as a fusion of the two chains (αβ)_2_ and the protein operates as a homodimer^23^. Only the α-chains resemble other Class II AARS in any way as they alone contain the three class-defining motifs, 1-3 defined by Eriani ^24^.

The GlyRS protein is from a distinct clade of the AARS phylogeny^3^ compared with our previous urzymes, which were from HisRS^10^, LeuRS^7, 25^, and TrpRS^11, 12^. For that reason, it is of particular interest. In this article, we demonstrate that the GlyCA urzyme shows strong aminoacylation activity. Much like our previous urzymes, the GlyCA is highly promiscuous, favoring amino acids activated by its own AARS Class (Class II). These results confirm that fully functional urzymes are readily obtained from wild type proteins and can even enter records as a result of contamination during genome assembly, therefore validating the robustness of the urzyme as an experimental model. We consider in detail here only the first 81 amino acids of ORF1. That fragment contains only motifs 1 and 2, previously shown sufficient for the HisCA2 urzyme^9, 10^ to aminoacylate tRNA^His^.

## Results

### Provenance, nomenclature, and likely architecture of the GlyCA urzyme

We identified the GlyRS urzyme sequence within the genome sequence for the Arctic Fox, *Vulpes lagopus*. We extracted the sequence from GenBank: gene number 121489478^26^. The annotation for the GenBank entry describes two open reading frames (ORFs 1,2; see Fig. 1). ORF-1 contains Motifs 1 and 2 plus 30 residues from the N-terminus of the insertion domain. These are the only structures homologous to other Class II AARS. ORF-2 contains a three-helix bundle from the C-terminus of the α-subunit. The GlyRS-B α-Chains assemble into dimers through that helical bundle^27^. These dimers can efficiently activate glycine even in the absence of the β subunit ^28^. They are not thought to have aminoacylation activity.

**Figure 1.**
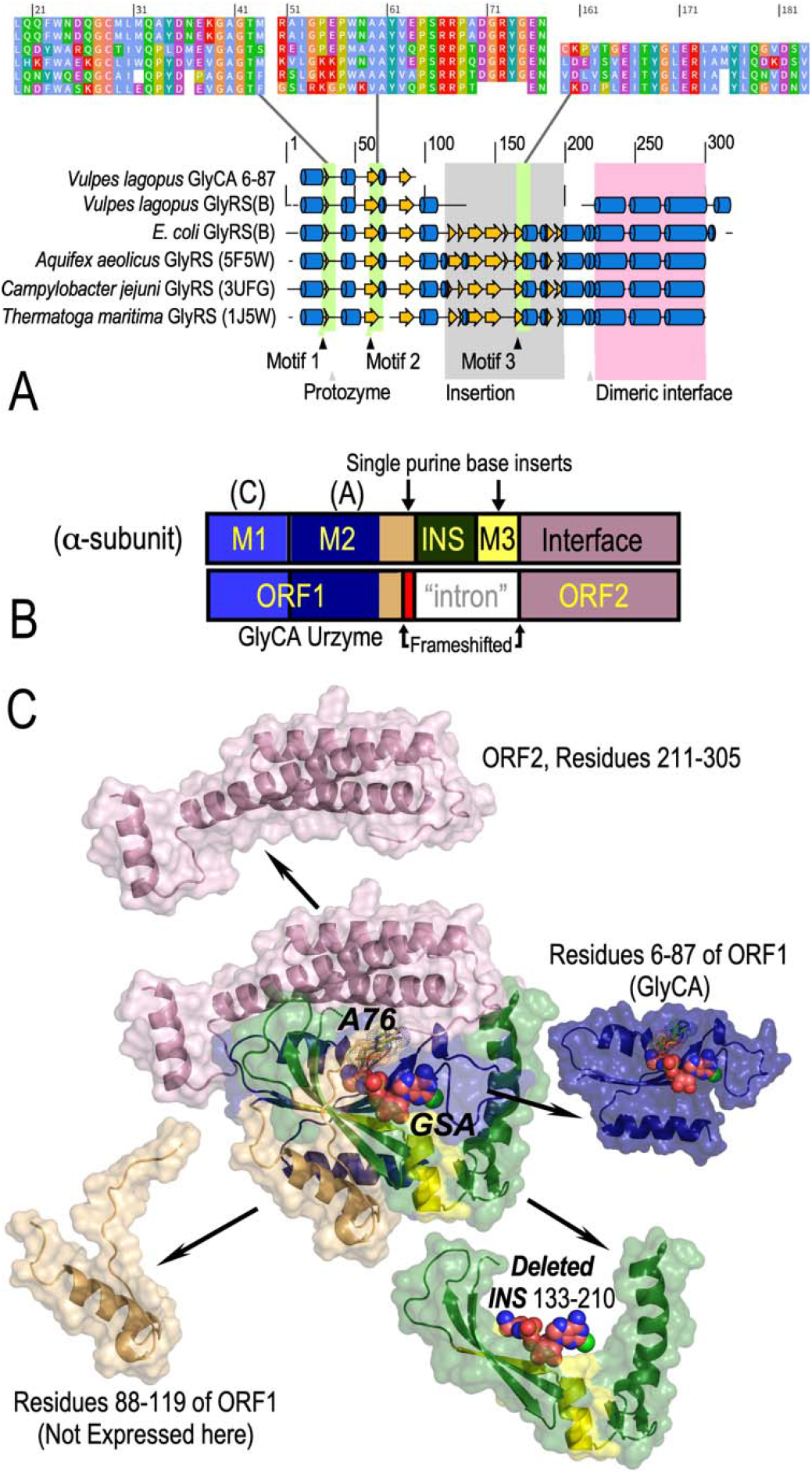
Architecture of the open reading frames from the *V. lagopus* GlyRS-B α-subunit. A. Schematic of unique secondary structures in heterotetrameric α_2_β_2_ GlyRS-B α-subunits. This figure shows a multiple structure alignment between the GlyCA AlphaFold structure and the alpha chains of three solved GlyRS-B structures (PDB codes indicated). Alpha helices are denoted by blue cylinders, beta strands by yellow arrows, while white space denotes a gap in the alignment. The alignment was generated by 3DCOMB^31^. Primary structures of Motifs 1, 2, and 3 that characterize Class II AARSs are shown above. The C-terminal alpha helix bundle is unique to GlyRS-B synthetases and forms the dimer interface. B. Altered modularity of the *V*. lagopus GlyRS α-chain, arising from 2 inserted purine bases (vertical arrows) that create frameshifting (tan) and an internal stop codon, producing two open reading frames. ORF1 ends with a stop codon C-terminal to the red frameshifted sequence. C. Alphafold2 prediction for the 3D structures of both open reading frames. Modules formed by the residue numbers indicated are exploded around the structure of ORFs 1,2. Colors are those used in B. AlphaFold2 prediction matches closely that observed in PDB ID 7YSE. That allows visualization of likely binding geometries for glycine-5’ sulfoamyl adenylate (GSA, spheres) and the 3’-terminal adenosine of tRNA^Gly^ (A76, dots). Motif 3 and an internal helix-turn-helix motif that covers the terminal adenosine in HisCA2^10^ are missing in GlyCA, leaving it with only 81 residues.

We recently introduced a nomenclature for Class I AARS urzymes, based on the sequential order of modules. The ATP-binding protozyme module containing the HIGH signature is N-terminal in all Class I AARS, and is denoted A. It also contains part of the amino acid binding site. A variable-length insertion element (connecting Peptide 1; CP1) follows the protozyme and contains various elements that enhance the amino acid specificity but are dispensable for both amino acid activation and aminoacylation. That insertion is segment B. The second signature, KMSKS, follows CP1^29, 30^. The anticodon-binding domain, which is always C-terminal in Class I AARS, is segment D.

We designated the LeuRS urzyme LeuAC because it contains segments A and C, but neither B nor D. Motifs 1 and 2 in Class II urzymes have high codon middle-base pairing frequencies in the opposite order in antiparallel alignment with Class I urzymes. Inasmuch as the Class II ATP-binding module (Motif 2) is C-terminal in Class II AARS, we will designate residues 6-87 of ORF1 as GlyCA (Fig. 1A).

We initially expressed purified two constructs as MBP-fusion proteins. The entire annotated gene (ORFs1+2) allowed us to determine the activity as represented directly in the genomic data. Residues 6-87 from ORF1 correspond to the Motif 1 and 2 segments of the simplest HisCA1 urzyme^10^. Active-site titration (not shown) revealed high burst sizes for both constructs, diagnostic for robust amino acid activation activity. We selected the 81-residue GlyCA for detailed studies.

Several features of that construct drew our attention.

- A BLAST search revealed that sequences of both open reading frames are 99.75% identical to the corresponding sequences of the bacterial GlyRS-B from *Streptococcus alactolyticus* sequenced from pig gut. Thus, it resembles a typical bacterial heterotetrameric GlyRS and likely arises either via contamination or, less likely, via horizontal gene transfer from a commensal bacterium.
- AlphaFold2 predicts a tertiary structure for the continuous coding sequences of ORFs 1 and 2 that is nearly identical to that of the catalytic center of the α-subunit in the 2.7 Å crystal structure of the *E. coli* α_2_β_2_ GlyRS-B^20^ (PDB ID 7YSE).
- There is no evidence of the mRNA being expressed in Arctic Fox tissues, and the splice site boundaries are computational predictions^26^.
- The premature stop codon is C-terminal to Motif 2, but N-terminal to the insertion module preceding Motif 3. It therefore includes the sequences homologous to the Class II HisCA urzymes^10^. We elaborate on this point in the DISCUSSION.
- The second open reading frame covers the entire C-terminal α-helical domain that forms the interface between the two α-subunits but is missing the insertion domain and Motif 3. Notably, GlyCA lacks the C-terminal three-helix bundle, so is expected to be monomeric.
- Notably, there is no sequence in the *V. lagopus* genome for a corresponding β subunit. The β-subunit, a highly idiosyncratic protein unlike any other AARS subunit, is absent. The β-subunit provides a variety of RNA binding domains and is normally required for aminoacylation.

### GlyCA is a stable, soluble, and functional aminoacyl-tRNA synthetase

We purified the GlyCA as an MBP fusion and characterized it directly, without removal of the tag by TEV cleavage. Purified GlyCA fusion protein is the primary source of observed catalytic activity. LC-ESI-MS/MS analysis (Fig. 2A) showed that GlyCA represented about 70% of the mass. The remaining proteins identified by the proteomics algorithm Protein Discoverer^32^ included numerous contaminants present at <3%. These contaminants included neither AARS nor other enzymes (kinases, etc.) expected to produce orthophosphate in the Malachite Green assay^33^. We confirmed that conclusion using a new zymographic technique^34^ (Fig. 2B) indicating that GlyCA was the only catalyst contributing to the achromatic green band in the zymogram gel. Single turnover kinetics show a high burst size (Fig. 2C). Previous studies of AARS urzymes relied heavily on active-site titration, AST—single turnover assays that estimate burst sizes—to demonstrate that observed catalytic activities originated from the principal component in purified samples^7, 10, 11, 25^. AST assays of GlyCA (Fig. 2C) show that nearly 90% of the urzyme molecules (0.88) contribute to ATP consumption. The nucleotide products of ATP consumption, ADP and AMP, occur in a 70:30 ratio. These data confirm the unique contribution made by the product translated from the GlyCA gene.

**Figure 2.**
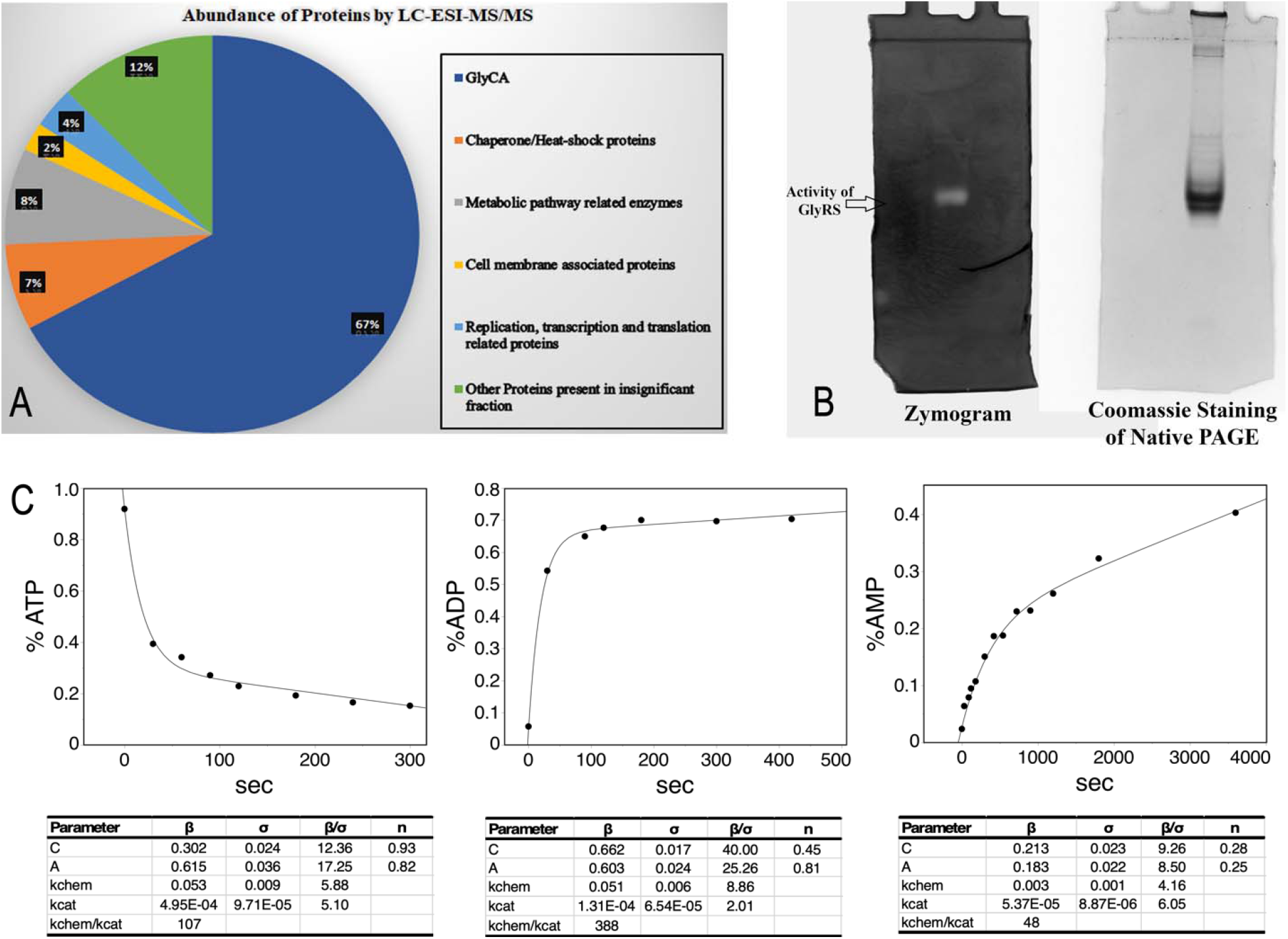
Purity and authentication of GlyCA catalysis. A. Mass spectroscopy of purified GlyCA. Liquid chromatography electrospray ionization mass spectroscopy was used to identify individual components of purified GlyCA. About 2/3 of the sample is GlyCA. The next 35 contaminants individually represent 0.03–0.002 of the sample and are broadly distributed among different metabolic pathways. None is capable of producing orthophosphate in the presence of amino acid and ATP. B. Comparison of a Coomassie stained native gel (right) to a zymogram visualized using absorption of the Malachite Green orthophosphate complex (left)^35^. C. Single turnover active site titration assay of GlyCA, showing fitted curves and fitting parameters for Eqn. 1 from densitometry of TLC plate autoradiographs. Use of ^32^Pα-ATP allows visualization of all three adenine nucleotides and thus the distribution of the two products, ADP and AMP. n-Values in the accompanying tables show that about 90% of the GlyCA molecules contribute to ATP consumption and that the two products, ADP and AMP, form with a ratio of ∼7:3^25^.

### GlyCA ORF1 catalyzes glycine activation

Michaelis-Menten kinetics of GlyCA [glycine]-dependence are shown in Fig. 3A, together with fitted kcat and K_M_ parameters. This and subsequent assays were done using the Malachite Green (MG) assay^35^. The picomolar sensitivity of the assay allows us to observe saturation behavior over an unusually large (30,000-fold) range of glycine concentrations (50 μM – 1.5 M).

**Figure 3.**
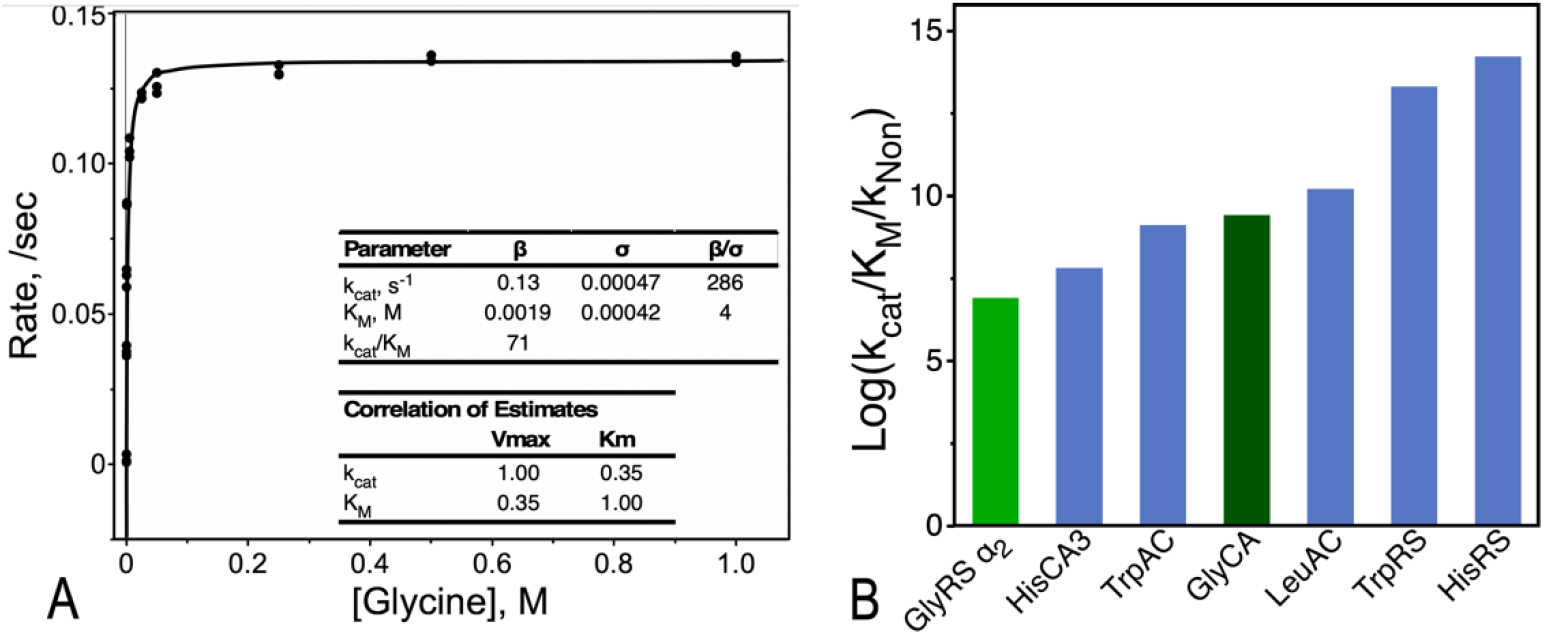
Michaelis-Menten kinetic analysis of glycine activation by GlyCA. **A**. Michaelis-Menten plot determined using the Malachite Green assay. The Michaelis-Menten rectangular hyperbola describes the glycine concentration dependence of triplicate rate measurements (R^2^ = 0.86) over a concentration range of 3 × 10^4^. K_cat_ and K_M_ parameter estimates are given with their variance parameters in the table. They are essentially uncorrelated. **B**. Comparison of the relative rate enhancements of GlyCA with other AARS constructs for amino acid activation. Note the substantial increase in GlyCA (dark green) over that of the intact GlyRS-B α_2_ dimer (light green).

Glycine binds relatively weakly, K_M_ = ∼2 mM, consistent with the absence of a side chain to provide either buried surface area or electrostatic attraction. Corresponding values for full-length tetrameric GlyRS-B enzymes (110 μM for *Aquifex aeolicus* ^*28*^; 160 μM for *E. coli* ^36^) are about 20-old tighter. The significant elevation of the GlyCA K_M_ over those of the two full-length GlyRS-B measurements reinforces our conclusion from Fig. 2 that the GlyCA urzyme is the sole source of observed catalytic activity.

The corresponding activation free energy for k_cat_/K_M_ = –2.5 kcal/mole, a value within the range (–3.6 to –1.3 kcal/mole) was previously established for other urzymes^2^. Notwithstanding the weak glycine binding, the overall rate enhancement, (k_cat_/K_M_)/k_Non_, (71/(8.3 × 10^−9^) = 8.5 × 10^9^)^12^ is nearly 10^10^-fold. That rate enhancement is ∼40 times greater than we previously observed for the HisCA3 urzyme^9^, and within a factor of 5 of that for LeuAC, the most active of the AARS urzymes characterized to date. Even more surprising, it is nearly 35 times greater than that measured for the intact *A. aeolicus* GlyRS-B α_2_ dimer^28^ (Fig. 3B).

### GlyCA catalysis of tRNA^Gly^ aminoacylation is comparable to that for Gly activation

GlyCA exhibits robust aminoacylation activity. The time-dependence measured with 115 μM tRNA^Gly^ and 4.7 μM GlyCA is shown in Fig. 4. The high tRNA^Gly^ concentration suggests that the fitted rate constant, 0.32 s^-1^, is close to k_cat_. That value is somewhat faster than comparable values for other urzymes. LeuAC aminoacylates tRNA^Leu^ with a k_cat_ = 0.088 s^-1^, but its high uncertainty suggests the values may be comparable^6^.

**Figure 4.**
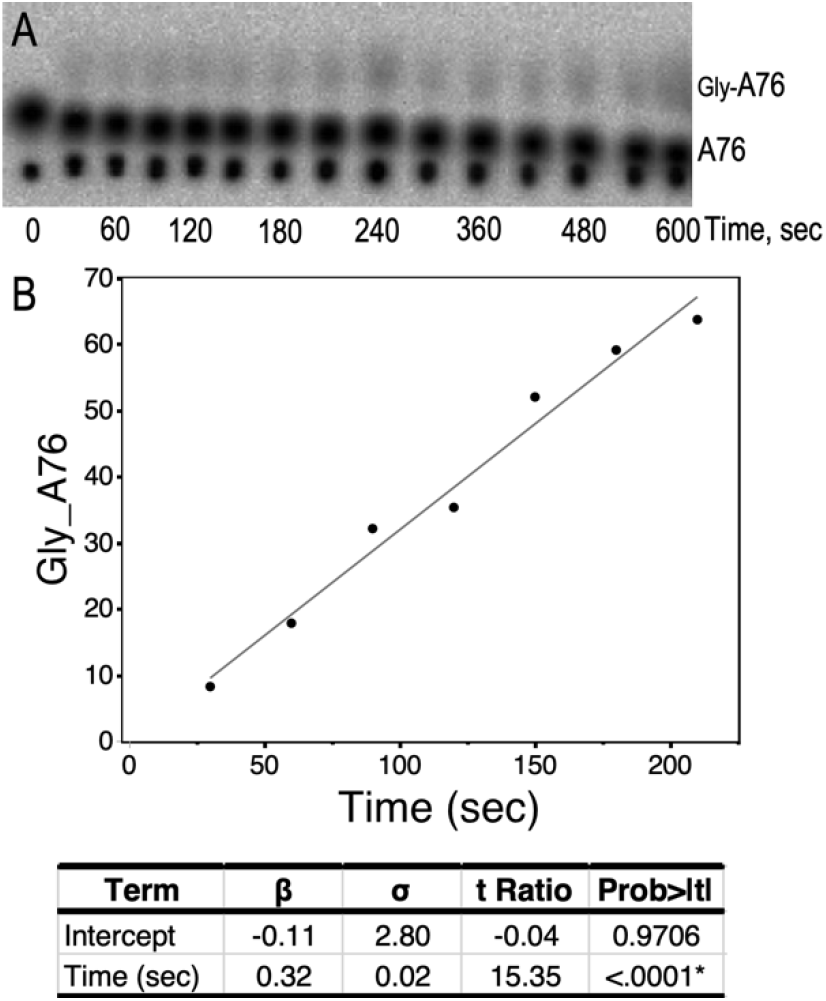
Time course of tRNA^Gly^ aminoacylation by GlyCA. **A**. Thin layer chromatogram showing time-dependent formation of Glycyl-tRNA^Gly^. **B**. The fitted linear portion of the time course.

Comparison with the activation rate constants provides additional perspective on the high k_cat_ value. It is about 3 times faster than the k_cat_ value for glycine activation (Fig. 3). The presence of tRNA enhances the rate of amino acid activation. We also observed a similar enhancement by TrpAC^9^, HisCA^9^, and LeuAC^6^ urzymes. Moreover, one of us (Tang, unpublished experiments) has observed that cognate tRNA increases first-order rate constants for the activation reaction in single turnover assays. Enhancement of amino acid activation rates by cognate tRNAs has often been described for full length AARS. Most notably, the Class IB GluRS, GlnRS, as well as LysRS and ArgRS cannot activate their cognate amino acids without their cognate tRNAs^1^. Discussions of coevolution of AARS with their cognate RNA substrates have appeared elsewhere^37^. Recent work has shown that the intimate functional cooperation between enzyme and RNA probably has roots in their most ancient functional ancestors^6^.

### GlyCA prefers Class II to Class I amino acids and recognizes a set of amino acids complementary to those recognized by other AARS urzymes

The Malachite Green assay enhances the throughput of enzymatic measurements, enabling us to perform Michaelis-Menten experiments for all 20 of the canonical amino acids. Free energies of activation for k_cat_/K_M_ are summarized in Fig. 5. The histogram divides the Class I (left) from Class II amino acids (right) and reflects the relative throughput of different amino acids and is therefore a quantitative metric for the ability of GlyCA to administer the code for glycine. It is worth noting that this specificity spectrum is the cleanest of the three we have published in the sense that glycine is preferred to a greater extent over other amino acids. Moreover, GlyCA favors a distinct set amino acids almost orthogonal to those favored by LeuAC and HisCA.

**Figure 5.**
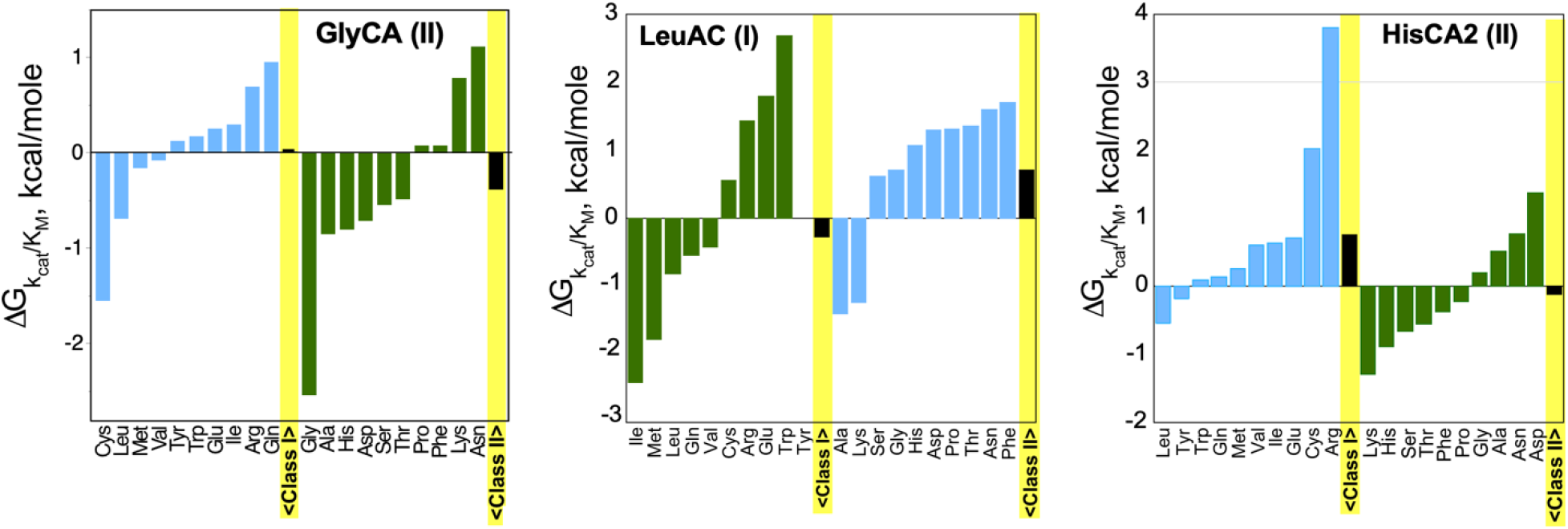
GlyCA amino acid specificity spectrum compared to those previously determined for LeuAC and HisCA2 (adapted from reference^2^). Michaelis-Menten experiments for GlyCA were performed using the Malachite Green assay as described in METHODS. Similar experiments for LeuAC and HisCA2 were performed using the conventional assay with radiolabeled ^32^P-pyrophosphate. The Class of each urzyme is appended in parentheses. Class I amino acids are sorted in the left, Class II amino acids on the right of each panel according to their activation free energies ΔG^‡^k_cat_/K_M_. Amino acids from the same Class as the urzyme are colored dark Green; those from the opposite Class are colored light blue. All three urzymes are promiscuous. GlyCA favors a subset of Class II amino acids (see yellow shaded columns showing Class averages in black) as does HisCA; LeuAC favors a subset of Class I amino acids. Amino acids favored by GlyCA (Gly, Ala) complement those favored by LeuAC (Ile, Met, Leu) and HisCA (Lys, His), suggesting a functional, experimental basis for implementing ancestral genetic codes with reduced alphabets.

## Discussion

### The GlyCA urzyme arose from a different paradigm from that of any previously characterized AARS urzymes

The GlyRS-B α-subunits are unique in retaining activity in the absence of β-subunits ^28^, which are themselves unique determinants of tRNA^Gly^ affinity. Identification of what appears to be a minimally modified *S. alactolyticus* <u>GlyRS-B</u> α-subunit is either a contaminant (*V. lagopus* is the only one of four fox genomes that have this gene) or the result of horizontal gene transfer.

The Arctic fox GlyCA and its matching gene (*Streptococcus alactolyticus* GlyRS) differ at two positions (pos 358 and 500). These two positions are 1nt insertions in the GlyCA sequence, which cause frameshift mutations (and therefore a premature stop codon). A single NT insertion after the codon of W119 creates a stop codon after residue D211, ending the first open reading frame (ORF1). The stop codon leads to “splicing” in the annotated database, and therefore the resulting protein sequence resembles GlyRS only at the N- and C-termini. The correct reading frame is restored downstream by the combination of a second inserted nucleotide and an intron that is not an integral00 multiple of codons in length. The annotated “intron” between ORF1 and ORF2 includes all of what is often referred to as the Insertion Domain, ID or INS ^38^, a three stranded antiparallel β module preceded by a short α-helix (Fig. 1C). Thus, derivation of the GlyCA architecture from its parent full-length GlyRS is entirely different from those of TrpAC, LeuAC, or HisCA urzymes, all of which were designed to test the Rodin-Ohno hypothesis ^18^. In that important sense, GlyCA is derived from observation, rather than by hypothesis.

The methods used for annotating intron-exon predictions are not the most accurate tools. They combine all kinds of information sources, not just identification of 5’ donor and 3’ acceptor splice sites in the putative pre-mRNA sequence. The identification of the *V lagopus* GlyRS-B 0ORFs 1 and 2 may be no more than coincidence, perhaps arising simply from sequencing errors. Notwithstanding, the GlyCA urzyme identified here proved to be a robust ancestral AARS construct.

### The rate acceleration for glycine activation, k_cat_/K_M_/k_Non_ = 8.6 s 10^9^, is ∼40-fold greater than the rate acceleration measured for HisCA3 (HisRS3 ^9^)

It is counterintuitive that this rate acceleration also exceeds that of the *A. aeolicus* α_2_-dimer by the same margin. One might expect that removing the helix bundle that provides the dimer interface might compromise enzymatic activity, rather than enhancing it. The authors describing the activity of the dimer note that it has a surprisingly reduced activity, relative to that of the α_2_β2 tetramer. It seems likely to us that this implies sophisticated communication between the two different subunits in the intact enzyme to coordinate catalysis of amino acid activation and tRNA acylation as well as specific recognition of both subunits. If that communication is mediated via the dimer interface, then it is possible that releasing the catalytic site from constraints imposed by the integrated behavior might activate it. Further experiments will be necessary for a fuller understanding.

### Evidence summarized in Fig. 2 represents the most extensive demonstration to date of the authenticity of urzyme catalysis

We supplement (i) the single turnover kinetic analysis (i.e., large burst size) and (ii) the two orders of magnitude increased K_M_ with (iii) the combination of LC_ESI_MS/MS and, especially (iv) zymographic evidence of *in situ* activity demonstrated in a native gel in documenting a comprehensive documentation of authenticity of the observed catalytic activity. These factors, together with the unusual provenance of GlyCA strongly underscores the legitimacy of using AARS urzymes as models for emergence and early evolution of the genetic coding table.

*Motif 3 is dispensable for rudimentary functionality of ancestral AARS in implementing a rudimentary genetic code*. We recently used thermodynamic cycle analysis to show that the active site signatures in Class I LeuAC also contribute only modestly to transition-state stabilization unless their functionalities are coupled together by the relative motions of domains not present in Class I urzymes^25, 39^. The absence of Motif 3 from the GlyCA urzyme reinforces our previous finding^9, 10^ that the second arginine tweezer^40^ is not required for catalysis of aminoacyl-tRNA synthesis. The ancestral requirements for enzymatic activity were surprisingly simple.

### AARS urzymes in the context of early genetic coding systems

Our work on AARS urzymes reveals several quantitative aspects of the kind of code they appear capable of implementing. We now have characterized three, phylogenetically diverse urzymes that share similar characteristics. They all exhibit a significant preference for amino acids from the Class of the amino acids that they administer as full-length contemporary enzymes. Further, although they lack the exquisite specificity of their mature forms, they each eliminate 4-6 non-overlapping amino acids from the ten within their own class. GlyCA for example prefers Gly, Ser, Ala, His, Asp, and Thr sufficiently for it to activate either Pro, Phe, Lys, or Asn less than 1time in ten.

As we have noted ^6, 41-43^, the coding of ancestral Class I AARS opposite and in-frame of ancestral Class II AARS provides structural bases for discriminating between both amino acid and tRNA acceptor stem substrates. Those rudimentary binding determinants act semi-independently as parallel filters, providing a basis for a rudimentary code. We also have noted ^44^ that an important barrier to enhancing the specificity may be a coding alphabet capable of discriminating between four redundant coding letters representing four mutually contrasting sets of amino acids.

This work opens a new perspective on the horizon, from which we may be able to frame a substantially clearer set of questions about the origin of the genetic code. Each of the AARS urzymes we have characterized behaves remarkably consistently with the requirements for a redundant four-letter code. The questions that now appear approachable will help us devise experiments with which to address whether or not such a code could be necessary and sufficient to create a foundry for protein evolution

## Methods

### The GlyCA urzyme sequence

The GlyAC sequence was extracted from GenBank: gene number 121489478^26^. We expressed both residues 6-87 of ORF1 and the fusion of ORFs1 and 2. Both have comparable activities. To avoid excessive complications due to putative quaternary structures arising from the presence of an extensive dimer interface, we consider in detail only the first exon: accession XM_041752554.1.

### Cell lysis and purification of GlyCA

Harvested cells were resuspended in a buffer containing 20 mM Tris, pH 7.4, 1 mM EDTA, 5 mM β-ME, 17.5% Glycerol, 0.1% NP40, 33 mM (NH_4_)_2_SO_4_, 1.25% Glycine, 300 mM Guanidine Hydrochloride plus cOmplete protease inhibitor (Roche). The cell suspension was lysed using a glass homogenizer followed by sonication (with 8 pulses of 10 second with 70% amplitude sonic vibrations) keeping 20 seconds of pause time, ensuring the tube remained in ice during sonication. GlyCA crude extract was then pelleted at 4 C with centrifugation of 15K rpm for 30 min to remove insoluble material. The extract supernatant was then diluted 1:4 with lysis buffer and loaded onto equilibrated Amylose FF resin (Cytiva). The resin was washed with five column volumes of buffer and the protein was eluted with 30 mM maltose in Optimal Buffer. The purified fraction was then dialyzed overnight with 50mM HEPES buffer containing 1 mM EDTA, 5 mM, 17.5% glycerol. After dialysis, fractions containing protein were pulled together, concentrated and mixed with 50% glycerol and stored at −80 C. Protein concentrations were determined using the Pierce™ Detergent-Compatible Bradford Assay Kit (Thermo Scientific)^7^.

### Label free proteomics; LC-ESI-MS/MS analysis

The protein lysates were first lyophilized and concentrated then dissolved in 50 mM Sodium acetate solution. 8M urea was added to the 150μg in-solution protein sample, then reduced with 5mM DTT for 30 min and alkylated with 15mM iodoacetamide for 45 min. The samples were diluted to 1M urea, then digested with MS grade trypsin (Promega) at 37 C overnight. Peptides were desalted with peptide desalting spin columns (Thermo) and dried via vacuum centrifugation. Each sample was analyzed in duplicate by LC-MS/MS using Easy nLC 1200 coupled to a QExactive HF (Thermo Scientific). Data analysis was done by Proteome Discoverer version 2.5 (Thermo Scientific).

The instrument used an Easy Spray PepMap C18 column (Thermo Scientific) and separated over a 120 min method. The gradient for separation was from 5 to 36% mobile phase B at a 250 nl/min flow rate, where mobile phase A was 0.1% formic acid in water and mobile phase B consisted of 80% acetonitrile, 0.1% formic acid. The QExactive HF was operated in data-dependent mode where the 15 most intense precursors were selected for subsequent HCD fragmentation. Resolution for the precursor scan (m/z 350–1700) was set to 60,000 with a target value of 3 × 10^6^ ions, 100ms inject time. MS/MS scans resolution was set to 15,000 with a target value of 1 × 10^5^ ions, 75ms inject time. The normalized collision energy was set to 27% for HCD, with an isolation window of 1.6 m/z. Peptide match was set to preferred, and precursors with unknown charge or a charge state of 1 and ≥ 8 were excluded.

Peptides detected by LC-ESI-MS/MS were matched with the *E. coli* database downloaded from Uniprot in FASTA format along with the GlyCA protein sequence. The following parameters were used to identify tryptic peptides: 10 ppm precursor ion mass tolerance; 0.02 Da product ion mass tolerance; up to 2 missed trypsin cleavage sites; (C) carbamidomethylation was set as a fixed modification; (M) oxidation was set as a variable modification. Peptide false discovery rates (FDR) were calculated by the Percolator node using a decoy database search and peptides were filtered using a 1% FDR cutoff^45^.

### Single turnover active-site titration assay of GlyCA

Active-site titration assays were performed using the same principle as described in Francklyn *et* al. and Fersht *et* al. ^46, 47^ with slight modifications. Briefly, 4 μM of GlyCA protein was added to a reaction mix containing 50 mM HEPES, pH 7.5, 10 mM MgCl_2_, 5 μM ATP, 50 mM Glycine, 1 mM DTT, inorganic pyrophosphatase, and α-labeled [^32^P] ATP to start the reaction. A volume of 2μL for all representative timepoints was added to separate tubes containing 4 μL quenching buffer (0.4 M sodium acetate, 0.1% SDS) and kept on ice until all time points had been collected. 3μL of Quenched samples were spotted on pre-run (PEI) TLC plates and run in TLC running buffer containing 850 mM Tris, pH 8.0. The plate was then dried and exposed for varying amounts of time to a phosphor image screen and visualized with a Typhoon Scanner (Cytiva). The intensities of each nucleotide were quantified by densitometry scanning using measure functions of ImageJ^48^. The time-dependences of loss (ATP) or de novo appearance (ADP, AMP) of the three adenine nucleotide phosphates were fitted using the nonlinear regression module of JMP™ Pro to Equation (1):

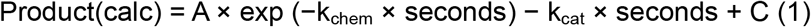

where k_chem_ is the first-order rate constant, k_cat_ is the turnover rate, A is the amplitude of the first-order process, and C is an offset.

### Zymography

To visualize amino acid activation in a native polyacrylamide gel chromogenically, Zymography was done using a 1.5 mm thick native gel of 8% resolving and 5% stacking which devoid of SDS^34^. Protein samples were prepared without adding SDS and β-ME to maintain the native conformation of protein^49^. Prepared protein (50μg) was then loaded into two separate wells of native gel. Electrophoresis was done with 40mA steady current at 4 °C and carefully observed. When the dye front reached the bottom of the gel, the gel was electrophoresed further for 30 minutes and then the electrophoresis was stopped. The gel was removed from the glass plates and placed in a glass box. Then the gel was washed with double distilled water for 5min in shaking condition and the step was repeated for two times.

After the washing step, substrate reaction buffer containing 50 mM HEPES of pH 7.5, 100mM glycine, 20mM MgCl_2_, 50mM KCl, pyrophosphatase solution [NEB] (0.1Unit/ml) and polyethylene glycol (PEG-8000; Sigma-Aldrich, Cat. No. 25322-68-3) into the mixture to a final concentration of 5% (w/v) was poured into the glass box containing the gel and keep the setup on a shaker for 45 minutes at 4°C. This step ensures complete soaking of substrate mixture into the gel; the low temperature reduces inactivation of the enzyme during the shaken perfusion.

After the perfusion most of the solution was decanted, leaving minimal solution in a static condition at 37°C. Amino acid activation was activated by adding 5mM ATP solution dropwise onto the gel surface to cover the whole gel. The gel was incubated in this condition for 30 minutes. After decanting the reaction mix from the gel box, the staining solution (0.05% Malachite green in 0.1 N HCl and 5% hexa-ammonium heptamolybdate tetrahydrate solution in 4 N HCl) was added directly onto the gel box. Staining was promoted by shaking for 2 minutes on a gyro-shaker.

The staining solution was made as described in Onodera *et* al.^33^. The gradual development of green bands (620 nm) around GlyCA protein present in the gel will signify the phosphomolybdate-malachite green complex formation and thus the *in situ* activity of amino acid activation by GlyCA. The gel was photographed using Gel Doc™ XR+ from BIO RAD imaging machine.

### Amino acid activation assays and Michaelis-Menten kinetics

Michaelis-Menten kinetics of amino acid activation assays for individual amino acids were done using a reaction mixture of 50mM HEPES of pH 7.0, 20mM MgCl_2_, 50mM KCl, 0.5mM ATP, 0.1 Unit of inorganic pyrophosphatase [NEB] and 7μM GlyCA enzyme and varied concentrations of a particular amino acid in separate tubes in a total reaction volume of 100 0000μL for each tube. The reaction was carried out at 37°C for 1 hour along with an enzyme blank in a separate tube. After this, 400 μL of malachite green-ammonium molybdate solution was added to the reaction tubes and kept for 5 minutes after mixing properly to develop the phosphomolybdate complex. 40uL of sodium citrate (w/v) were then added to the tubes and then solutions were allowed to stand for 20 min and the optical absorbance at 620nm was measured with DU800 spectrophotometer (Beckman Coulter Inc., Brea, CA, USA). The malachite green-ammonium molybdate solution was prepared as described by Onodera *et*. al.,^33^. The phosphoric acid concentration was calculated from a calibration curve prepared by using 0–250 μM K_2_HPO_4_, which indicates the relationship between phosphoric acid concentration and absorbance. The specific activity of the GlyCA for individual amino acid concentrations were calculated and plotted in a specialized non-linear fit using the Michalis-Menten equation in JMP™ Pro software. The K_M_ and k_cat_ of GlyCA for individual amino acids were determined, with standard deviations, from the maximum likelihood fit.

## Acknowledgements

A major part of the LC-ESI-MS/MS analysis and data processing is conducted using the UNC Proteomics Core Facility, which is supported in part by P30 CA016086 Cancer Center Core Support Grant to the UNC Lineberger Comprehensive Cancer Center.

## Author contributions

S.K.P. expressed, purified and assayed GlyCA, and performed all experimental measurements, including zymography, AST, aminoacylation, and sample preparation for LC-MS-MS analysis.

L.B. designed the expression vector. J.D. identified the gene as a possible Class II AARS urzyme and provided phylogenetic insight for Figure 1. P.W. and R.B. C.W.C validated the analyses. C.W.C., S.K.P. and J.D. wrote the manuscript, All authors contributed to discussions throughout the course of the work, and all contributed and approved the final figures and text.

## Data availability

All data will be provided upon request from the author.

---

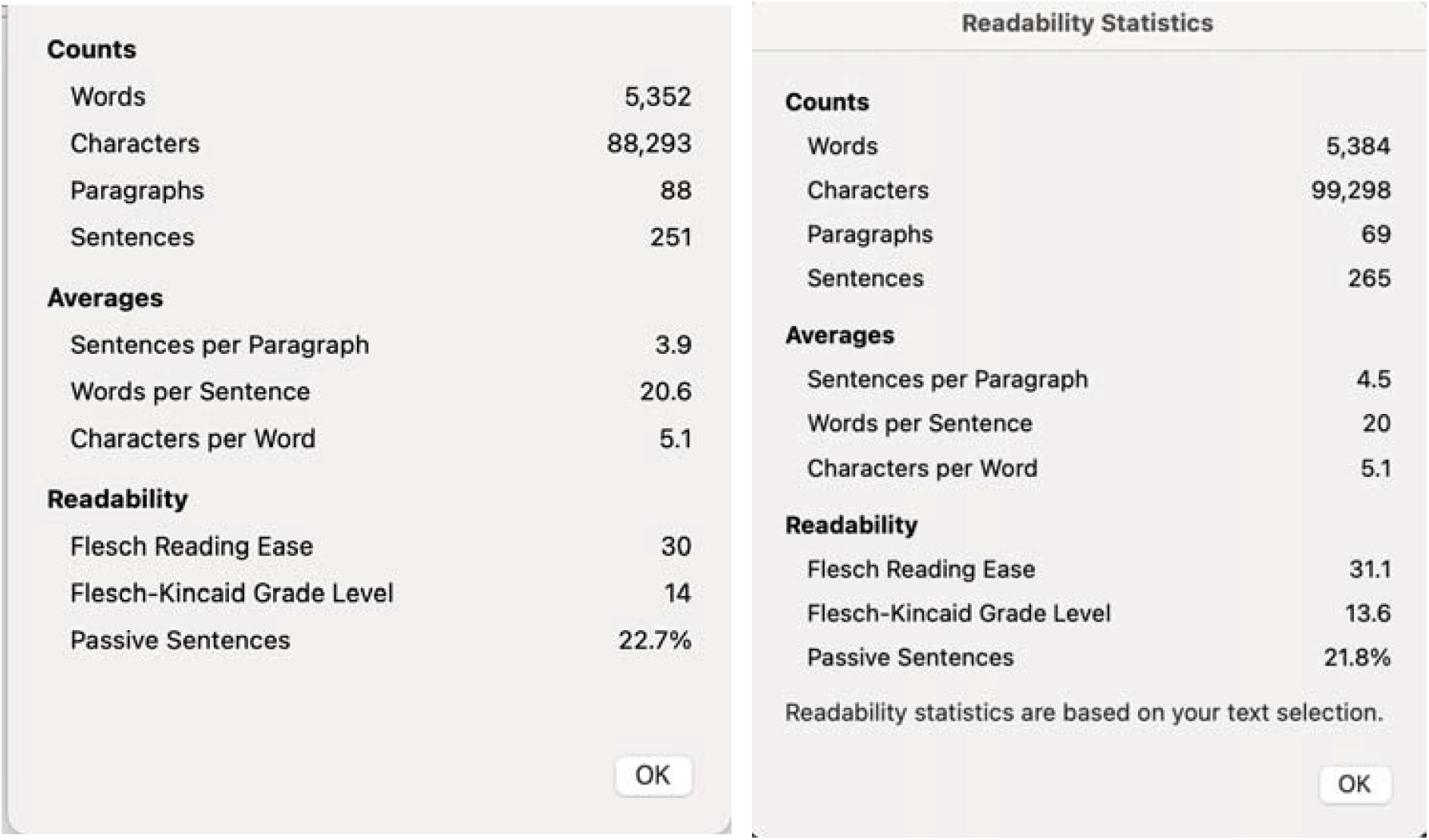

